# Detection and identification of bacterial *spp.* from culture negative surgical site infection patients using molecular tool

**DOI:** 10.1101/689992

**Authors:** Himanshu Sekhar Behera, Nirupama Chayani, Madhusmita Bal, Sanghamitra Pati, Sashibhusan Das, Hemant Kumar Khuntia, Manoranjan Ranjit

## Abstract

Surgical site infection (SSI) is the most common post operative infections, that sometimes cause postoperative morbidity, mortality and increase in hospital costs. Managing surgical site infections, with negative culture report in routine diagnosis is a common dilemma in microbiology. The present study was carried out to know the presence and frequencies of various bacteria in wound aspirates/ swabs of some culture negative surgical site infection patients attending a tertiary care hospital of eastern India with molecular methods. Ninety seven (97) patients with post-operative SSI whose wound swabs/ aspirate were negative in the conventional aerobic culture after 48 hrs of incubation were taken for finding the presence and identification of any bacteria in sample with 16S rRNA gene specific broad PCR and submitted to NCBI. Of the 97 patients, 16S rRNA based broad range PCR assay could able to identify the presence of bacterial pathogen in 53(54.63%) cases, out of which 29 isolates were of viable but non-culturable bacteria(VBNC), 07 were of obligatory anaerobes and 13 were of unculturable bacteria, 04 were with polybacterial infections. Some measures should be taken in the microbiology laboratory along with conventional culture, for better patient care such as, culture plates should be allowed to incubate for an additional 3 – 4 days that will allow the growth any fastidious bacteria, anaerobic culture system should be developed along with aerobic system, and molecular diagnosis by 16S broad range PCR should be performed routinely.

## Introduction

Maintaining and improving the quality in healthcare service is a major concern in hospitals and other health care facility. Surgical site infections (SSI) are the most common post operative infections of skin or underlying soft tissue that sometimes causes postoperative morbidity, mortality, increase hospital costs and prolongs hospital stay(1, 2). SSI is the third most commonly reported nosocomial infections after ICU infections and urinary tract infections (UTI) in a hospital set up, approaching ~16% of all nosocomial infections(3). As per Centre for Disease Control and Prevention (CDC) and the European Centre for Disease Prevention and Control (ECDC)), SSI is defined as, “postoperative infection occurring within 30 days of surgery or within one year if any prosthetic material is implanted at the surgical site”(4). There are some ways by which surgical sites can get infected, such as: use of unsterile instruments, contaminated prosthetics and even surgical solutions while performing surgical procedures; improper cleaning of surgical site by infected surgical solutions which allows the entering of skin flora such as *Staphylococcus epidermidis*, *Mycoplasma* species etc into the surgical site, which later causes infections(5). Sometimes some pathogens such as *Mycoplasma* species present in the blood reach the surgical site and causes SSI(2). It is recognized by “purulent discharge around the wound or the insertion site of the drain, or spreading cellulitis from the wound” (1, 5). SSI varied from 2.5% to as high as 41.9% based on hygienic conditions of clinical set ups(6).

Culture negative surgical site infection is a common problem while furnishing a report in a microbiology laboratory, which is defined as “a patient with all the clinical signs of surgical site infection, but with “no bacterial growth” in the conventional culture(2). Incidence of such ‘culture negative SSIs’, reported in some of the studies can be up to 30%(7). There are several causes of culture negative SSI, such as: if the sample is collected from the patient after commencement of antibiotics, delayed SSI, presence of fastidious, viable but non-culturable bacteria(VBNC) or unculturable bacteria in sample etc(2). In all these cases culture plate will be sterile even after incubation for 72hrs. In some cases, though there might be the presence of very small colonies of bacteria such as: Atypical *Mycobacteria, Mycoplasma and Ureaplasma, Legionella*, Small-colony variant” *Staphylococcus aureus* in plate, they are either missed or over looked by the microbiologist, hence sterile culture report get furnished to patient(2, 5). Sometimes due to the presence of common contaminant like *Staph. epidermidis* in the culture plate, they are often discarded, but they may be actually responsible for post operative SSI(2). As only aerobic culture system is available in most of the microbiology laboratories, the anaerobic bacteria present in sample cannot grow in that aerobic conditions(2). Sometimes the culture media or growth conditions are not supportive for the growth of some bacteria, hence no bacterial growth is noticed even after the incubation of plate for 72 hrs(8). Those unculturable bacteria must be growing in their natural environment, but we are not able to grow them in laboratory conditions, still they sometimes cause infections. Some studies had also reported the presence of several unculturable bacteria in clinical specimen(9).

Conventional culture is the Gold standard to identify the causative pathogen in clinical samples, but the results are completely dependent on the presence of viable organism and time of processing of samples after collection(10). If the culture plate is sterile even after incubation of plate for 48 or 72hrs, it does not means that, there is always no organism in the sample(10). Therefore there is always a need for a broad spectrum, rapid diagnostic method to detect and identify the causative bacteria, when the culture plate is sterile even after incubation for 48/ 72 hrs. After the invention of PCR, molecular diagnosis was attempted with PCR for rapid and accurate diagnosis of all diseases. The 16S rRNA gene is present in all bacteria, which comprised of many genus specific or species specific conserved and variable regions. Broad-range 16S rRNA gene specific PCR assay is useful in identifying the causative pathogen even in multiple bacterial infections, presence of fastidious organisms in sample, patient with prior anti microbial therapy or presence of any unculturable bacteria in sample etc(11). The present study was carried out to know the presence and frequencies of various bacteria in aspirates/ wound swabs of some culture negative surgical site infection patients attending a tertiary care hospital of Odisha, an eastern India State using molecular tools.

## Materials and Methods

### Selection of patients and collection of clinical specimens

In this prospective study, 97 patients with post-operative Surgical Site Infections (SSI) referred from the surgery department of SCB Medical College, Cuttack from March 2018 to February 2019, whose swabs/ wound aspirate were negative in the conventional culture were included in this study. Of the 97 patients 32 were females and 65 were males, having their age from 28 yrs to 84yrs. Surgical site infection patients whose clinical samples (swabs/ aspirates) were positive in conventional culture after 48 hrs of incubation were excluded from this study. Ethical clearance was obtained from human ethical committee of the institute and samples were collected after informed consent of the patient. Wound aspirates and swabs were collected from the Microbiology department of SCB Medical College, Cuttack after the sample comes sterile with conventional culture and processed for molecular analysis using 16S rDNA based broad range PCR assay.

### Molecular diagnosis

#### DNA isolation

DNA was extracted from the pus/ wound aspirate specimens using commercial QIAamp DNA Mini Kit (Qiagen), as per the manufacturer’s instructions.

#### Broad-range PCR assay

Briefly, broad-range PCR assay was standardized to amplify ~1492 bp region of 16S rRNA gene using published primers (FP: (27F) 5’-AGAGTTTGATCCTGGCTCAG-3’ and RP: (1492R) 5’-GGTTACCTTGTTACGACTT-3’) while varying the annealing temperature and conc. of MgCl_2_(12). PCR amplification was carried out in 25 μl of final reaction volume containing 1× reaction buffer (Fermentas), 0.2 mM dNTPs (Fermentas), 0.40 μM of each primer (IDT) and 1.25U Taq polymerase (Fermentas). The temperature profile of the PCR assay was as follows: initial denaturation for 04 mins at 94°C, followed by 35 cycles of denaturation for 1min at 94°C, primer annealing for 1min at 57°C, strand elongation for 1min at 72°C, with the final elongation for 10mins at 72°C temperature. DNA isolated from known isolates of *E. coli* was used as a positive control and reaction mixture with 5μl of distilled water was used as a negative control in all PCR reactions. Amplified PCR products were electrophoresed on 1.0 % agarose gel and visualized under a gel documentation system (Syngene).

#### Nucleotide sequencing and homology analysis

Amplified DNA bands from 16S rRNA gene based PCR assay were cut with sterile scalpel blades from the agarose gel and processed for Sanger nucleotide sequencing (Eurofins). The obtained nucleotide sequences were searched for homology analysis with the available sequences of 16S rRNA gene of the GenBank using NCBI BLAST (NCBI, USA) computer program (http://www.ncbi.nlm.nih.gov/pubmed). Nucleotide sequences of samples which showed >90% homology with the available 16S rRNA sequences of any bacteria were submitted to NCBI to obtain the respective accession numbers.

## Results

Of the ninety seven (97) culture negative surgical site Infection (SSI) patients, 16S rRNA based broad range PCR assay could able identify the presence of bacterial pathogen in 53(54.63%) cases, all of which were successfully sequenced through Sanger sequencing. Of the 53 nucleotide sequences, (n=12; 22.2%) were found belongs to *Bacillus* spp., (n=13; 24.07%) were uncultured bacterium, (n=06; 11.1%) *Pseudomonas* spp., (n=06; 11.1%) were of *Enterococcus* Spp., (n=02; 3.7%) were *Bacteroides*, (n= 02; 03.7%) were *Fussobactrium* sp., (n=02; 3.7%) *Massilia* sp., (n=01; 01.8%) *Staphylococcus* sp., (n=01; 1.8%) *Sneathia* sp., (n=1; 1.8%) *Peptostreptococcus* spp., (n=1; 1.8%) *Kleibsiella* sp., (n=1; 1.8%) *Stenotrophomonas* sp., (n=1; 1.8%) *Peptoniphilus* sp., (n=4; 7.4%) samples were of multiple bacterial infections(as per Sanger sequencing results). Of the 53 samples positive for PCR assay, 49 were submitted to NCBI data bank and obtained respective accession numbers, which are given in Table 1.

**Table 1.** Accession numbers of the bacteria identified and submitted to NCBI.

| **Serial No.** | **Sample name** | **Bacteria identified** | **Accession No.** | **Gram positive (GP)/ Gram Negative (GN)** | **Obligate anaerobic(OA )/ Facultative anaerobic(FA)** |
| --- | --- | --- | --- | --- | --- |
| 1 | SSI 1 | Bacillus spp. | MK424355 | GP | FA |
| 2 | SSI 2 | Bacillus spp. | MK424356 | GP | FA |
| 3 | SSI 4 | Staphylococcus spp. | MK424357 | GP | FA |
| 4 | SSI 5 | Sneathia spp. | MK424358 | GN | OA |
| 5 | SSI 6 | Uncultured bacterium | MK424359 |  |  |
| 6 | SSI 7 | Bacillus spp. | MK424360 | GP | FA |
| 7 | SSI 9 | Mixed infections | NA |  |  |
| 8 | SSI 10 | Mixed infections | NA |  |  |
| 9 | SSI 11 | Pseudomonas spp. | MK424361 | GN | FA |
| 10 | SSI 12 | Bacillus spp. | MK424362 | GP | FA |
| 11 | SSI 13 | Mixed infections | NA |  |  |
| 12 | SSI 14 | Uncultured bacterium spp. | MK928502 |  |  |
| 13 | SSI 15 | Bacteroides | MK928506 | GN | OA |
| 14 | SSI 16 | Fussobacterium spp. | MK928505 | GN | OA |
| 15 | SSI 18 | Fussobacterium spp. | MK928504 | GN | OA |
| 16 | SSI 20 | Uncultured bacterium spp. | MK424363 |  |  |
| 17 | SSI 21 | Bacteroides | MK928503 | GN | OA |
| 18 | SSI 22 | No significant similarity | NA |  |  |
| 19 | SSI 24 | Bacillus spp. | MK424364/ | GP | FA |
| 20 | SSI 26 | Mixed infections | NA |  |  |
| 21 | SSI 27 | Enterococcus spp. | MK424365 |  |  |
| 22 | SSI 28 | Pseudomonas spp. | MK424366 | GN | FA |
| 23 | SSI 29 | Bacillus spp. | MK829054 | GP | FA |
| 24 | SSI 31 | Enterococcus spp. | MK829055 | GP | FA |
| 25 | SSI 32 | Massilia spp. | MK838102 | GN | FA |
| 26 | SSI 34 | Uncultured bacterium spp. | MK934354 |  |  |
| 27 | SSI 35 | Uncultured bacterium spp. | MK424367 |  |  |
| 28 | SSI 38 | Uncultured bacterium spp. | MK858273 |  |  |
| 29 | SSI 39 | Uncultured bacterium spp. | MK424368 |  |  |
| 30 | SSI 40 | Uncultured bacterium spp. | MK829056 |  |  |
| 31 | SSI 41 | Peptostreptococcus spp. | MK858274 | GP | OA |
| 32 | SSI 42 | Uncultured bacterium sp. | MK934355 |  |  |
| 33 | SSI 44 | Bacillus spp. | MK424369 | GP | FA |
| 34 | SSI 45 | Bacillus spp. | MK858266 | GP | FA |
| 35 | SSI 46 | Enterococcus spp. | MK829057 | GP | FA |
| 36 | SSI 48 | Enterobacter spp. | MK424370 | GP | FA |
| 37 | SSI 49 | Enterobacter spp. | MK424371 | GP | FA |
| 38 | SSI 50 | Massilia spp. | MK858267 | GN | FA |
| 39 | SSI 51 | Klebsiella spp. | MK424372 | GN | FA |
| 40 | SSI 57 | Pseudomonas spp. | MK838103 | GN | FA |
| 41 | SSI 60 | Peptoniphillus spp. | MK928501 | GP | OA |
| 42 | SSI 61 | Stenotrophomas spp. | MK838104 | GN | FA |
| 43 | SSI 64 | Uncultured bacterium sp. | MK934356 |  |  |
| 44 | SSI 66 | Bacillus spp. | MK829058 | GP | FA |
| 45 | SSI 68 | Pseudomonas spp. | MK829059 | GN | FA |
| 46 | SSI 71 | Pseudomonas spp. | MK829060 | GN | FA |
| 47 | SSI 73 | Uncultured bacterium spp. | MK934357 |  |  |
| 48 | SSI 76 | Uncultured bacterium spp. | MK838105 |  |  |
| 49 | SSI 80 | Pseudomonas spp. | MK858268 | GN | FA |
| 50 | SSI83 | Bacillus spp. | MK858269 | GP | FA |
| 51 | SSI 87 | Bacillus spp. | MK858270 | GP | FA |
| 52 | SSI 91 | Uncultured bacterium<br>spp. | MK928500 |  |  |
| 53 | SSI 93 | Bacillus spp. | MK858271 | GP | FA |
| 54 | SSI 97 | Enterococcus spp. | MK858272 | GP | FA |

Of the 49 identified bacteria, 29 isolates were of viable but non-culturable(VBNC) bacteria belonging to *Bacillus* spp., *Pseudomonas* spp., *Enterococcus* spp., *Massilia* spp., *Staphylococcus* spp., *Kleibsiella* spp., *Stenotrophomonas* spp. and 07 were of obligatory anaerobic strains belonging to *Bacteroides* spp., *Fussobactrium* spp., *Sneathia* spp., *Peptostreptococcus* spp.*, Peptoniphilus* spp. and 13 were of unculturable strains. Out of 49 bacterial isolates identified in this study 21 belongs to Gram-positive bacteria and 15 belongs to Gram-negative bacteria. Of all the bacteria 03 were of slow growing (fastidious) nature that are of genus *Massilia* spp. and *Peptostreptococcus* spp. respectively.

## Discussions

Managing surgical site infections, with negative culture report in routine diagnosis is a common problem in the microbiology laboratory. The frequency of SSI depends upon the type of surgery performed and the hospital environment. Similarly prevalence of pathogens in SSI varies from place to place and hospital to hospital(13). Studies reported that, around 5–30% of clinical specimens (wound swabs/ aspirates) isolated from a patient having all clinical signs of SSI, do not show any bacterial growth in conventional culture(6). In a study conducted in a tertiary care hospital in southern India (Bangalore) 7.8% of all SSI were culture negative(14). Similary in a study conducted in a medical college in western India (Maharashtra), out of 196 pus samples taken from SSI patients 5.4% were negative in culture(15). Similarly other studies conducted across India and World have reported several cases of culture negative SSI. In those cases, although there is the presence of bacteria in some samples, other factors plays a crucial role in preventing their growth on culture plate such as, sample collection after the commencement of antibiotics, presence of viable but non-culturable bacteria(VBNC) and fastidious bacteria in the sample, if the laboratory condition is not suitable for the growth of anaerobic bacteria and the presence of unculturable bacteria in samples etc (2, 9).

In this study we have reported several facultative aerobic culturable bacteria such as *Bacillus* spp., *Pseudomonas* spp., *Enterococcus* spp., *Massilia* spp., *Staphylococcus* spp., *Kleibsiella* spp., *Stenotrophomonas* spp., from wound swabs/ aspirate of culture negative SSI patients, which points towards their viable but non-culturable(VBNC) state, that results the culture plate turn negative after an incubation of 48hours. VBNC bacteria refers to the bacteria which remain in a state of very low metabolic activity, do not divide but are alive and does not appear in conventional culture. As this is a tertiary care hospital, most of the patients come here after long term antibiotic treatment from primary health care hospitals, hence most of the pathogenic bacteria transformed into viable but non-culturable(VBNC) state due to either stress, biofilm or spore formation. In one of the review article, all these bacterial species are reported to transform to VBNC state in stress conditions(16). Many research aticles have reported the presence of these VBNC bacteria in patient samples, which can be identified by PCR assay(17, 18). Some studies also reported that, pathogenic bacteria such as *Bacillus* sp., *Pseudomonas* sp., *Staphylococcus* sp. turn to slow or non growing bacterial cells called persisters to survive from lethal dose of antibiotics, hence they did not come in culture plate within 48 hrs of incubation(19, 20). Hence our study corroborates with the previous studies.

Most of the SSI due to anaerobic bacteria are derived from the host’s own endogenous flora, with few exceptions like *Clostridium* spp. These endogenous anaerobic bacteria play a vital role in preventing the colonization of several pathogenic and exogenous microbial populations, but due to some structural or functional defects in the mucous layer or obstructions become pathogenic(21). The predominant anaerobic bacteria which are involved in SSI include *Bacteroides fragilis* group, *Prevotella* spp., *Porphyromonas* spp., *Fusobacterium* spp., *Peptostreptococcus* spp, *Clostridium* spp. and *Actinomyces* spp.(22). In this study, we have reported the presence of some anaerobic bacteria such as; *Bacteroides*, *Fussobactrium* spp., *Sneathia* spp., *Peptoniphilus* spp. and *Peptostreptococcus* spp. in the wound swab/ aspirates of culture negative SSI patients. As there is no anaerobic culture set up available in this medical college, even though anaerobic bacteria was present in some samples, they cannot grow on culture plate in aerobic conditions(22). Hence the diagnostic laboratories in medical colleges must set up both aerobic and anaerobic microbial culture facility to support the growth of both aerobic and anaerobic bacteria in clinical samples that will be beneficial for the patient care.

Unculturable bacteria was also reported in 13 patient samples out of 97 culture negative SSI samples. Currently the developed culture media is not able to grow them in laboratory conditions, but in near future some culture media will be able to support the growth of these unculturable bacteria. Therefore attention should be made to develop suitable laboratory media/ conditions to support the growth of these currently uncultured bacteria (10). This is a great challenge but an opportunity to work in this area. For this, it is essential to understand the metabolism of these bacteria.

Managing a patient with no growth in culture after 48hrs of incubation is a challenging task. Certain experimental measures can be taken to improve the diagnosis of the patient if the sample is culture negative after 48hrs of incubation. In those cases culture plates should be allowed to incubate for an additional 3 – 4 days, which will allow the growth of any fastidious bacteria if present in sample. As anaerobic culture system is rarely available in the microbiology set up in India, it should be developed, so that any anaerobic bacteria if present in sample should be identified in culture. As several unculturable bacteria are also responsible of culture negative SSI, molecular diagnosis by 16S broad range PCR assay is quite useful in identifying their presence in sample. This will help the clinicians in prescribing appropriate antibiotic to the patient. As molecular assay has more sensitivity and accuracy in identifying the pathogen than culture, this should be employed in routine diagnosis.

## Acknowledgements

Authors thank all the clinical faculty members and resident doctors of SCB medical college for sending the clinical specimens for clinical investigations.

## Conflicts of interest satements

The authors declare that, there are no conflicts of interest related to this work.

